# Type I Interferon Transcriptional Network Regulates Expression of Coinhibitory Receptors in Human T cells

**DOI:** 10.1101/2020.10.30.362947

**Authors:** Tomokazu S. Sumida, Shai Dulberg, Jonas Schupp, Helen A. Stillwell, Pierre-Paul Axisa, Michela Comi, Matthew Lincoln, Avraham Unterman, Naftali Kaminski, Asaf Madi, Vijay K. Kuchroo, David A. Hafler

## Abstract

While inhibition of T cell co-inhibitory receptors has revolutionized cancer therapy, the mechanisms governing their expression on human T cells have not been elucidated. Type 1 interferon (IFN-I) modulates T cell immunity in viral infection, autoimmunity, and cancer, and may facilitate induction of T cell exhaustion in chronic viral infection^1,2^. Here we show that IFN-I regulates co-inhibitory receptors expression on human T cells, inducing PD-1/TIM-3/LAG-3 while surprisingly inhibiting TIGIT expression. High-temporal-resolution mRNA profiling of IFN-I responses enabled the construction of dynamic transcriptional regulatory networks uncovering three temporal transcriptional waves. Perturbation of key transcription factors on human primary T cells revealed both canonical and non-canonical IFN-I transcriptional regulators, and identified unique regulators that control expression of co-inhibitory receptors. To provide direct *in vivo* evidence for the role of IFN-I on co-inhibitory receptors, we then performed single cell RNA-sequencing in subjects infected with SARS-CoV-2, where viral load was strongly associated with T cell IFN-I signatures. We found that the dynamic IFN-I response *in vitro* closely mirrored T cell features with acute IFN-I linked viral infection, with high *LAG3* and decreased *TIGIT* expression. Finally, our gene regulatory network identified *SP140* as a key regulator for differential *LAG3* and *TIGIT* expression. The construction of co-inhibitory regulatory networks induced by IFN-I with identification of unique transcription factors controlling their expression may provide targets for enhancement of immunotherapy in cancer, infectious diseases, and autoimmunity.

## Main

Immune checkpoint blockade targeting T cell co-inhibitory receptors has revolutionized cancer treatment. While signals such as IL-27 are associated with T cell expression of co-inhibitory receptors, in mice^3,4^, the regulatory mechanisms of co-inhibitory receptors induction in human T cells are still unknown. Type 1 interferon (IFN-I) are induced during chronic viral infection, autoimmunity, and cancer^5–7^, and accumulating evidence suggests that IFN-I may have immunomodulatory function beyond their conventional role of promoting generation of effector T cells^8–11^. Notably, continuous exposure to IFN-I is implicated to promote T cell exhaustion, which is marked by aberrant expression of co-inhibitory receptors (i.e. PD-1, TIM-3, LAG-3, and TIGIT) in chronic viral infection and cancer^11–16^. However, whether IFN-I facilitates induction of immune checkpoint molecules and T cell function, directly or indirectly, has not been investigated.

## The impact of IFN-β on co-inhibitory receptors in human T cells

We have previously identified IL-27 as a crucial cytokine that promotes the induction of a co-inhibitory receptor module, with T cell exhaustion in a murine tumor models^3^. Several reports have suggested that IL-27 functions downstream of IFN-β^17^, which is a major component of the IFN-I family. Thus, we hypothesized that IFN-β facilitates the induction of co-inhibitory receptors in humans. We first assessed the effect of IL-27 and IFN-β on induction of core co-inhibitory receptors (TIM-3, LAG-3, PD-1, and TIGIT) *in vitro* using human primary naïve CD4^+^ and CD8^+^ T cells (Supplementary Figure 1a). Both IL-27 and IFN-β promoted significantly higher TIM-3 expression compared to control condition without addition of exogenous cytokines. Of note, IFN-β induced more LAG-3 and TIM-3 expression compared to IL-27 in both CD4^+^ and CD8^+^ T cells (Figure 1a, b, Supplementary Figure 1b). Unexpectedly, both IL-27 and IFN-β suppressed the expression of TIGIT in CD4^+^ and CD8^+^ T cells (Supplementary Figure 1c). We also observed the increased production of IL-10 by IFN-β; however, IL-10 induction by IL-27 was modest in our *in vitro* culture settings, which may reflect the difference between mouse and human T cell responses toward IL-27 stimulation (Supplementary Figure 1d). To determine whether these observations stemmed from the effect of IFN-β on cellular proliferation, we performed a proliferation assay using cell trace violet dye. We found there was no differences in cellular division in naïve CD4^+^ T cells between control and the IFN-β condition. Additionally, there was even less proliferation of memory CD4^+^ T cells in the IFN-β condition, indicating that the induction of co-inhibitory receptors by IFN-β is not driven by a state of higher proliferation in T cells (Supplementary Figure 2), consistent with previous studies^18,19^. We further determined the impact of IFN-β treatment on gene expression kinetics for co-inhibitory receptors by qPCR. Gene expression dynamics for core co-inhibitory receptors *(HAVCR2, LAG3, PDCD1)* were upregulated by IFN-β for most time points. In contrast *TIGIT* was downregulated, which was confirmed by protein expression using flow cytometry (Figure 1c, d). We examined the expression of other co-inhibitory receptors, and found IFN-β induced co-expression of multiple co-inhibitory receptors (e.g. *HAVCR2, PDCD1, LAG3),* but inhibited expression of others *(TIGIT, CD160, BTLA)* (Figure 1e). Collectively, these data elucidate a role for IFN-β as a cytokine that can directly control multiple co-inhibitory receptors in both human CD4^+^ and CD8^+^ T cells *in vitro.*

**Figure 1.**
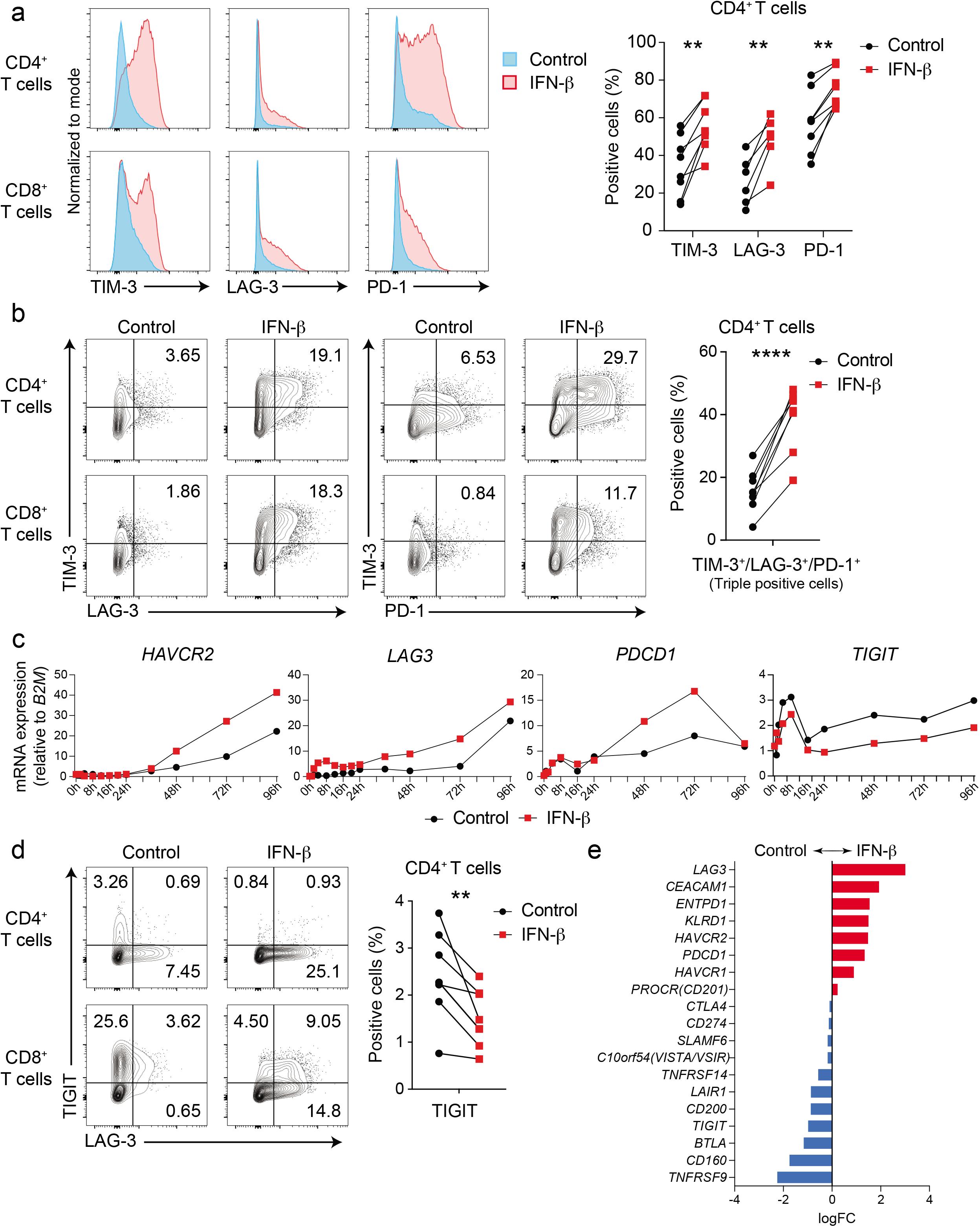
IFN-β differently regulates LAG-3, TIM-3, PD-1 and TIGIT in human T cells. Effects of IFN-β on LAG-3, TIM-3, PD-1, and TIGIT expression on human naïve CD4^+^ and CD8^+^ T cells cultured with anti-CD3/CD28 for 96h in the absence (Control) or with 500 U/ml IFN-β (IFN-β). **a,** Representative histograms of flow cytometry analysis (left), quantitative expression for LAG-3, TIM-3, and PD-1 expression on naïve CD4^+^ T cells (n = 6 – 8) (right). **b,** Representative contour plots of flow cytometry analysis on surface LAG-3, TIM-3, and PD-1 (left), quantitative analysis for LAG-3, TIM-3, and PD-1 triple-positive cells in naïve CD4^+^ T cells (n = 8) (right). **c,** Gene expression kinetics of *LAG3, HAVCR2, PDCD1,* and *TIGIT*quantified by qPCR with 13 timepoints in naïve CD4^+^ T cells. Average expression values from two subjects are plotted. **d,** IFN-β induces LAG-3 but suppresses TIGIT expression on human naïve CD4^+^ and CD8^+^ T cells. Representative contour plots of flow cytometry analysis (left), quantitative analysis for TIGIT positive cells in naïve CD4^+^ T cells (n = 8) (right). **e,** Co-inhibitory receptors expression pattern under IFN-β treatment in naïve CD4^+^ T cells by qPCR (n = 4). Red and blue bars represent higher expression in IFN-β treatment and Control condition, respectively. Data was represented as mean +/− SD. **p < 0.01, ****p < 0.0001. Paired Student’s t test.

## Transcriptomic dynamics of IFN-β response at high temporal-resolution

To uncover the regulatory mechanisms underlying the IFN-β response in human primary T cells, we generated a transcriptional profile at high temporal resolution. We used bulk mRNA-seq at ten time points along a 96-hour time course with and without IFN-β treatment (Supplementary Figure 3a). To avoid inter-individual variation, we selected one healthy subject whose T cells exhibited a stable response to IFN-β, and repeated the experiment three times at a two-week interval for each experiment. We identified 1,831 (for CD4^+^ T cells) and 1,571 (for CD8^+^ T cells) differentially expressed genes (DEGs) across time points with IFN-β treatment, revealing a temporal shift of gene expression patterns in both CD4^+^ and CD8^+^ T cells (Figure 2a, Supplementary Figure 3b, Methods). The genome-wide transcriptional profiles from three independent experiments demonstrated highly replicative results across time points (Figure 2b). Specifically, three transcriptional waves were observed during 96 hours of IFN-β response: early phase (1-2h), intermediate phase (4-16h), and late phase (48-96h). As expected, we observed abundant induction of classical IFN stimulated genes (ISGs) (i.e., *IFI6, MX1/2, RSAD2, STAT1/2/3, SP100/110/140),* which peaked at the early-intermediate phase. IFN-I induced cytokines produced by T cells *(IFNG, IL10, GZMB, PRF1)* were also upregulated at intermediate-late phase (Figure 2c, Supplementary Figure 3c). Interestingly, OSM, which is reported to amplify IFN-β response and suppress Th17 differentiation^20^, was significantly induced by IFN-β from the early phase, and maintained induction in all time points (Supplementary Figure 3c). Among DEGs, we identified dynamic expression of 134 TFs for CD4^+^ T cells and 100 TFs for CD8^+^ T cells, which were both up- and down-regulated over the course of differentiation (Figure 2c).

**Figure 2.**
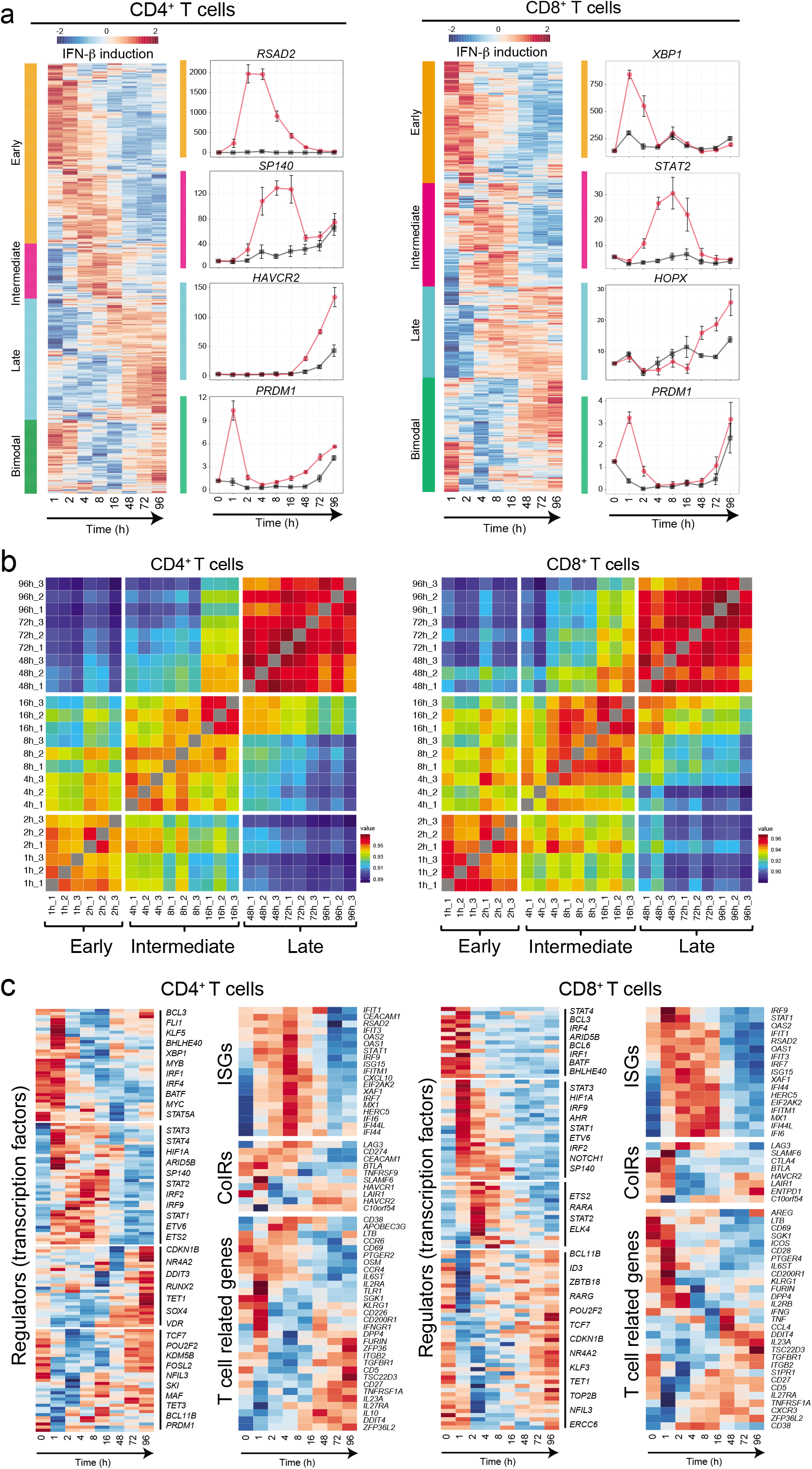
Three waves of dynamic transcriptomic changes by IFN-β in human T cells. **a,** Gene expression profiles under IFN-β treatment in naïve CD4^+^ and CD8^+^ T cells. Differential expression of gene levels for eight time points with IFN-β stimulation (log2(expression)) are shown in heatmap. Based on the expression kinetics, the genes are clustered into four categories: early, intermediate, late, and bimodal (up regulated at early and late phase). Representative individual gene expression kinetics from each cluster are shown (mean+/− SD). **b,** Correlation matrix of global gene expression representing three transcriptional waves on CD4^+^ (left) and CD8^+^ (right) T cells: early (1-2h), intermediate (4-16h), and late (48-96h). Eight timepoints with three replicates are shown. **c,** Temporal transcriptional profiles of differentially expressed genes for four categories are shown; transcriptional regulators (transcription factors), ISGs, co-inhibitory receptors, and key T cell associated factors for CD4^+^ (left) and CD8^+^ (right) T cells.

## Lentiviral shRNA based genetic perturbation

To narrow the list of TFs for perturbation, we prioritized TFs that are differentially expressed in both CD4^+^ and CD8^+^ T cells. We then further selected TFs associated with: 1) human tumor infiltrating T cells (TILs)^21–24^; 2) HIV specific T cell signature in progressive patients^25^; and 3) IL-27 driven co-inhibitory regulators^3^ (see Methods). We confirmed that these TFs are identified as interferon stimulated genes (ISGs) in human immune cells by the Interferome dataset^26^ (Figure 3a). In total, 31 TFs were listed as candidates based on the overlap between ISG, TIL and IL-27 signatures and we chose 19 of them for perturbation. Since TCF-1 (encoded by *TCF7)* and Blimp-1 (encoded by *PRDM1)* are known to express functionally distinct isoforms, we also targeted a unique sequence for the long isoforms *(TCF7L* and *PRDM1L,* respectively), resulting in perturbation of 21 different targets in total.

**Figure 3.**
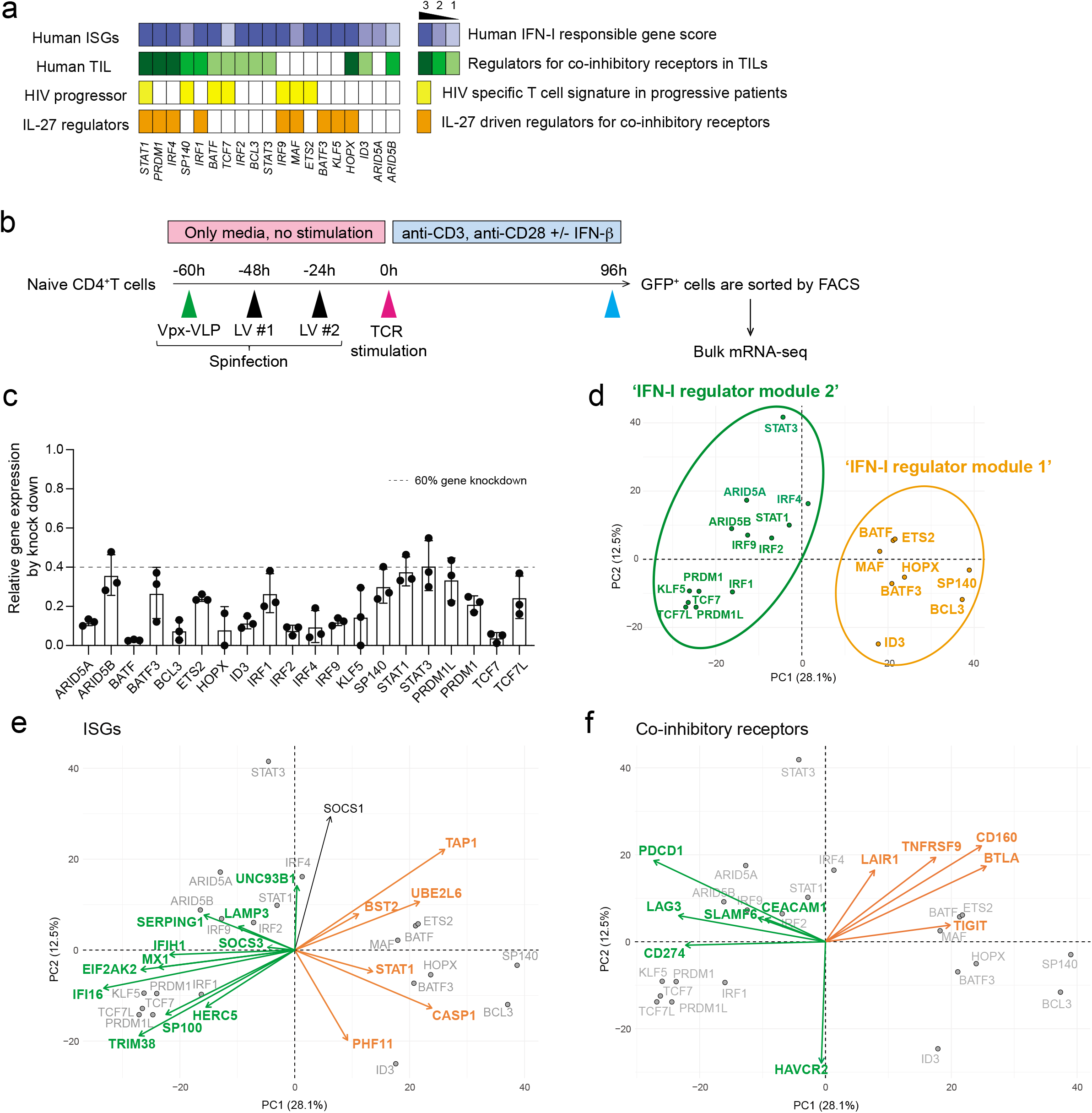
Perturbation of key transcription factors in quiescent human T cells. **a,** Characterization of candidate TFs for perturbation. Perturbed TFs are listed based on overlap between differentially expressed TFs of CD4^+^ T cells and CD8^+^ T cells. Human ISG score (top; blue), human TIL co-inhibitory receptors score (green), HIV specific T cell signature genes in progressive patients (yellow), and IL-27 driven co-inhibitory receptor regulators (orange) are shown for each TFs. **b,** Experimental workflow of Vpx-VLP supported lentiviral shRNA perturbation. Ex vivo isolated naïve CD4 T cells were transduced with Vpx-VLPs, followed by two times of lentiviral particle transduction before starting T cell activation. T cells were stimulated with anti-CD3/CD28 in the absence or presence of IFN-β (500 U/ml) for 96h and GFP positive cells were sorted by FACS. RNAs were extracted from sorted cells and applied for mRNA-seq. Perturbation for all 21 shRNAs are performed with human CD4^+^ T cells isolated from the same individual as in Figure 2. **c,** Gene knockdown efficiency is shown as relative expression over scramble shRNA transduced controls. Dotted line represents 60% of gene knockdown. **d-f,** PCA plots and biplots based on differentially expressed genes by perturbation. **d,** PCA plot demonstrating the two modules of TF regulators on perturbation with 21 TFs. Characterization of shRNA-based gene knockdown for each TF being plotted. Labels represent perturbed TF gene names. ‘IFN-I regulator module 1’ is colored in green and ‘IFN-I regulator module 2’ is in orange. **e, f,** PCA biplot showing differential regulation by modules of regulator TFs; **e,** for ISGs and **f,** for co-inhibitory receptors. Orange and green arrows (vectors) are highlighting two groups of genes effected inversely by the different modules of TFs.

Considering the majority of these regulators are induced at the early (1-2h) and intermediate (4-16h) phases, it was important to perform gene deletion prior to T cell receptor activation. For human primary T cells gene knockdown, we adopted lentiviral delivery of shRNAs with lentiviral gene product X (Vpx) containing virus-like particles (VLPs) system to efficiently transduce lentivirus into unstimulated primary human naïve T cells^27,28^ (Figure 3b). Spinoculation with Vpx-VLPs significantly increased the number of GFP expressing T cells (~30-60%) compared to normal LV particle transduction without Vpx-VLPs (~1-5%), and resulted in successful transduction of lentiviral vectors into non-blasting/quiescent cells (Supplementary Figure 4a). We achieved efficient knockdown of at least 60% gene expression for 21 target TFs in human naïve CD4^+^ T cells (Figure 3c).

To identify the effect of perturbation for each regulator, Principal Component Analysis (PCA) was applied to changes in RNA expression associated with each transcription factor knock down (Figure 3d). PC1 divided the impact of perturbation into two modules of regulators; *BATF, MAF, ETS2, HOPX, SP140, BCL3, ID3,* and *BATF3* constitute ‘IFN-I regulator module 1’, and *IRF1, IRF2, IRF4, STAT1, STAT3, ARID5A, ARID5B, TCF7, PRDM1, PRDM1L, KLF5, and TCF7L* constitute a distinct ‘IFN-I regulator module 2’. To visualize the contribution of the selected genes to the PCs and the directionality of the contribution, a PCA biplot analysis was adopted. We found that ISGs are divided into two groups; classical ISGs that are correlated with ‘IFN-I regulator module 1’ (depicted in green arrows in Figure 3e), and ISGs that are correlated with ‘IFN-I regulator module 2’ (depicted in orange arrows in Figure 3e), which is predominantly in PC1. These results suggest that ISGs are bi-directionally regulated by different modules of TFs (Supplementary Figure 4b). Furthermore, these modules contributed differently to the regulation of co-inhibitory receptors; *TIGIT, CD160, BTLA* as one module and *LAG3, HAVCR2, PDCD1* as another module (Figure 3f, Supplementary Figure 4c). Within ‘IFN-I regulator module 1’, STAT3 positively regulated *HAVCR2* but not *LAG3* and *PDCD1* expression, which is predominantly contributed by PC2 in the biplot (Figure 3f). Taken together, our perturbation with Vpx-VLP system elucidated two distinct TF modules simultaneously regulating opposing IFN-I target genes, which sheds light on the central roles of non-canonical IFN-I induced regulators in human T cells.

## Dynamic transcriptional regulatory network of IFN-β response

To characterize the impact of differentially expressed transcription factors (DETFs) in response to IFN-β, we generated transcriptional regulatory networks describing TFs and their target genes for each of the transcriptional waves identified (Figure 4a, Methods). When comparing the three regulatory networks, early and late networks had similar numbers of TFs (46 and 42 TFs, respectively), while the intermediate network contained 73 TFs (Figure 4b, top). Interestingly, the ratio between up and down-regulated TFs differs between the three regulatory waves. The early and intermediate network contained more up-regulated TFs than down-regulated TFs; in contrast, the late network had more down-regulated than up-regulated TFs. Thus, IFN-I induced differentiation involves dominance of up-regulated TFs in the first 16 hours, replaced by the dominance of down-regulated TFs after 48 hours. We next ranked the TFs based on the enrichment of their target genes and their centrality in the networks (Methods), highlighting the significance of each TF to the network (Figure 4b, middle). In the early regulatory network, *MYC* and *CDKN1B/KDM5B* were among the most dominant up and down-regulated TFs, respectively, in this transcriptional wave. These data indicate that T cell metabolic activation, cell cycle regulation, and transcriptional activation are promoted by IFN-β treatment^29,30^. Interestingly, *FLI1,* which is novel as a IFN-I downstream TF and was recently reported to control effector response in T cells^31^, also dominantly regulated the early transcriptional wave. In the intermediate regulatory network, *MYC*, *MAF, IRF1, AFF1, ATF3, and TBX21* were among the most dominant up-regulated TFs for this transcriptional wave. In the late network, effector function related regulators are upregulated *(PRDM1, RUNX2, MAF, BCL3);* in contrast, the TFs associated with Treg differentiation and maintenance *(STAT5A, FOXP1, MYB)* were down-regulated, suggesting the skewed differentiation toward effector-like signature.

**Figure 4.**
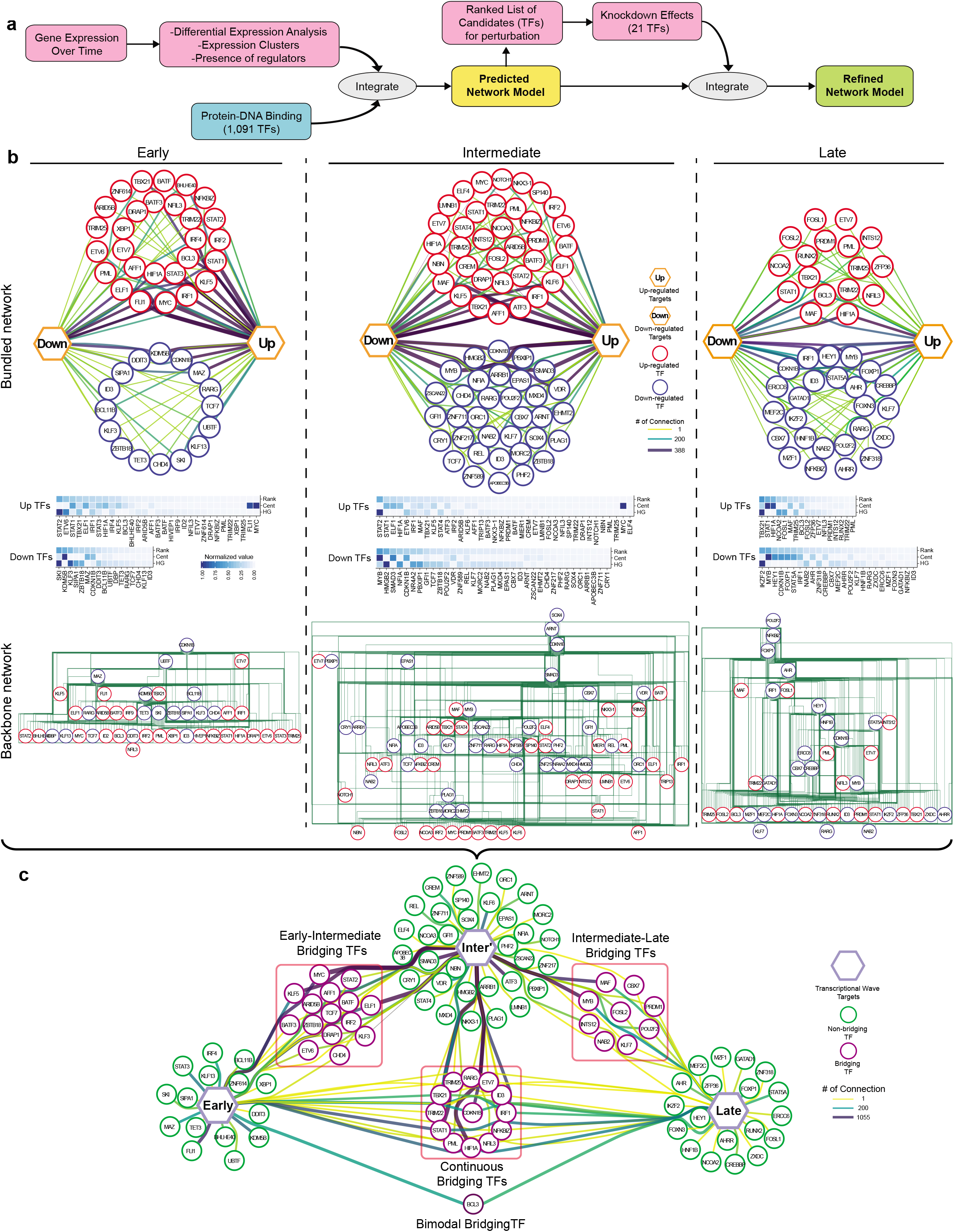
Transcriptional regulatory network under IFN-I response. **a,** Overview of regulatory network generation. A scheme of the pipeline, in order to generate preliminary regulatory network is generated from integrating the gene expression kinetics data coupled with TF-target gene datasets. The key regulators’ perturbation data was further integrated to refine the preliminary network. **b,** In depth view of the transcriptional regulation at each wave. Top row; the representation of regulatory networks highlighting TFs interaction. The thicker and darker an edge is the more TF-target connections it represents. Target genes are represented by up and down hexagons, according to their regulatory response to IFN-β. Middle row; heatmaps representing a ranking of the TFs based on ‘Cent’ stands for centrality and ‘HG’ stands for hypergeometric test. Bottom row; hierarchical backbone networks. Red circles represent up-regulated TFs, blue circles represent down-regulated TFs. **c,** Dynamics of TFs regulation across the transcriptional waves. Each hexagon represents targets from each transcriptional wave. Green circles represent regulatory TFs which are differentially expressed only in one transcriptional wave they are connected to, while purple circles represent bridging TFs, which are DE in all transcriptional waves they are connected to. The thicker and darker an edge is, the more TF-target connections it represents.

To further study the relationships between the DETFs, we generated hierarchical backbone networks in order to represent their relationships (Figure 4b, bottom). Interestingly, the top TFs in all transcriptional time waves were down-regulated in response to IFN-β, while TFs lower in the network hierarchy were more up-regulated. A few examples from the early transcriptional wave hierarchical backbone include *CDKN1B* and *MAZ*, which appeared at the top of the hierarchy, whereas *KLF5, MYC,* and *FLI1* were lower in the hierarchy. It was of interest that these TFs were also highly dominant in the regulatory network. These data suggest that loss of suppression of TFs at higher hierarchy triggers the activation of downstream effector TFs under IFN-I response, which was also observed in the intermediate and late regulatory network. The elucidation of this backbone network enables us to shed light on the regulatory interactions within each component of the transcriptional network, providing further depth to the extent of interactions within the network.

While T cell differentiation under IFN-β is characterized by three major transcriptional waves, we hypothesized that there are key TFs that bridge each wave to the next. To this end, we specifically identified TFs that participate in more than one of these transcriptional waves, and termed these ‘Bridging TFs’ (Figure 4c). Examples of dominant ‘Bridging TFs’ between early and intermediate waves include *KLF5* and *STAT2.* Examples of intermediate to late waves include *MAF, PRDM1,* and *MYB.* Finally, there are TFs that were upregulated throughout the entire differentiation; such as *STAT1, HIF1A,* and *TBX21.* Generally, ‘Bridging TFs’ tend to be more dominant than other TFs; thus, it is possible that ‘Bridging TFs’ play an important role in the transition between different transcriptional waves. Indeed, our perturbation experiment demonstrated the critical roles of those ‘Bridging TFs’ in the regulation of ISGs and co-inhibitory receptors (Figure 4c). Our computational analysis revealed the temporal dynamics of complex regulatory interactions during the IFN-I response and highlighted the usefulness of our approach in discovering this new aspect of IFN-I induced transcriptional regulation.

## *In vivo* validation of regulatory modules controlling IFN-I/co-inhibitory receptors axis in human T cells

As it was important to provide direct *in vivo* evidence for the role of IFN-I on T cell co-inhibitory receptor expression, we sought to validate our regulatory network in the human setting where the IFN-I response of T cells is induced acutely. As acute viral infections are strongly associated with IFN-I responses, we examined a number of clinical models where viral infection is closely linked to IFN-I T cell response. We found that our analysis of single cell RNA seq (scRNA-seq) analysis of T cells in COVID-19 patients revealed an extremely high correlation between viral load and IFN-I score (r=0.8) and time difference between paired samples and the respective change in IFN-I score (r=0.97)^32^, providing a unique opportunity to generate a rich dataset to determine whether the *in vitro* T cell response to IFN-I can be validated during an acute viral human infection strongly associated with a IFN-I signal.

By using our scRNA-seq data, we subclustered T cell populations into 13 subpopulations and identified five CD4^+^ T cell and five CD8^+^ T cell subsets (Figure 5a, Supplementary Figure 5a, b). We first focused on total CD4^+^ T cells and CD8^+^ T cells and confirmed that the IFN-I response signature is higher in progressive patients who required admission to the ICU and eventually succumbed to the disease (Figure 5b). Expression of co-inhibitory receptors differed across disease conditions, but the trend was conserved between CD4^+^ and CD8^+^ T cells. We observed a strikingly similar pattern of co-inhibitory receptor expression with IFN-I stimulation *in vitro* and *in vivo.* Indeed, we observed the upregulation of ‘IFN-I up co-inhibitory receptors’ *(LAG3/HAVCR2)* and the downregulation of ‘IFN-I down co-inhibitory receptors’ *(TIGIT/LAIR1/SLAMF6)* in T cells from COVID-19 patients (Figure 5c, d). As expected, ‘IFN-I up co-inhibitory receptors’ were positively correlated with canonical ISGs expression, but ‘IFN-I down co-inhibitory receptors’ were not, suggesting that there are different regulatory mechanisms dictating co-inhibitory receptor expression patterns (Figure 5e). Next, we investigated which subpopulation of CD4^+^ and CD8^+^ T cells is more affected by the IFN-I response, and computed the IFN-I score across subpopulations for each T cell subtype in COVID-19 patients (Figure 5f). Within the subpopulations that exhibited higher IFN-I scores, dividing CD4^+^/CD8^+^ T cells and ISG^+^ CD8^+^ T cells were uniquely increased in COVID-19 patients, particularly in severe patients^32,36^. Moreover, these subpopulations and effector T cells expressed higher level of co-inhibitory receptors and ‘IFN-I regulator module-1’ compared to the other subpopulations (Figure 5g, Supplementary Figure 5c).

**Figure 5.**
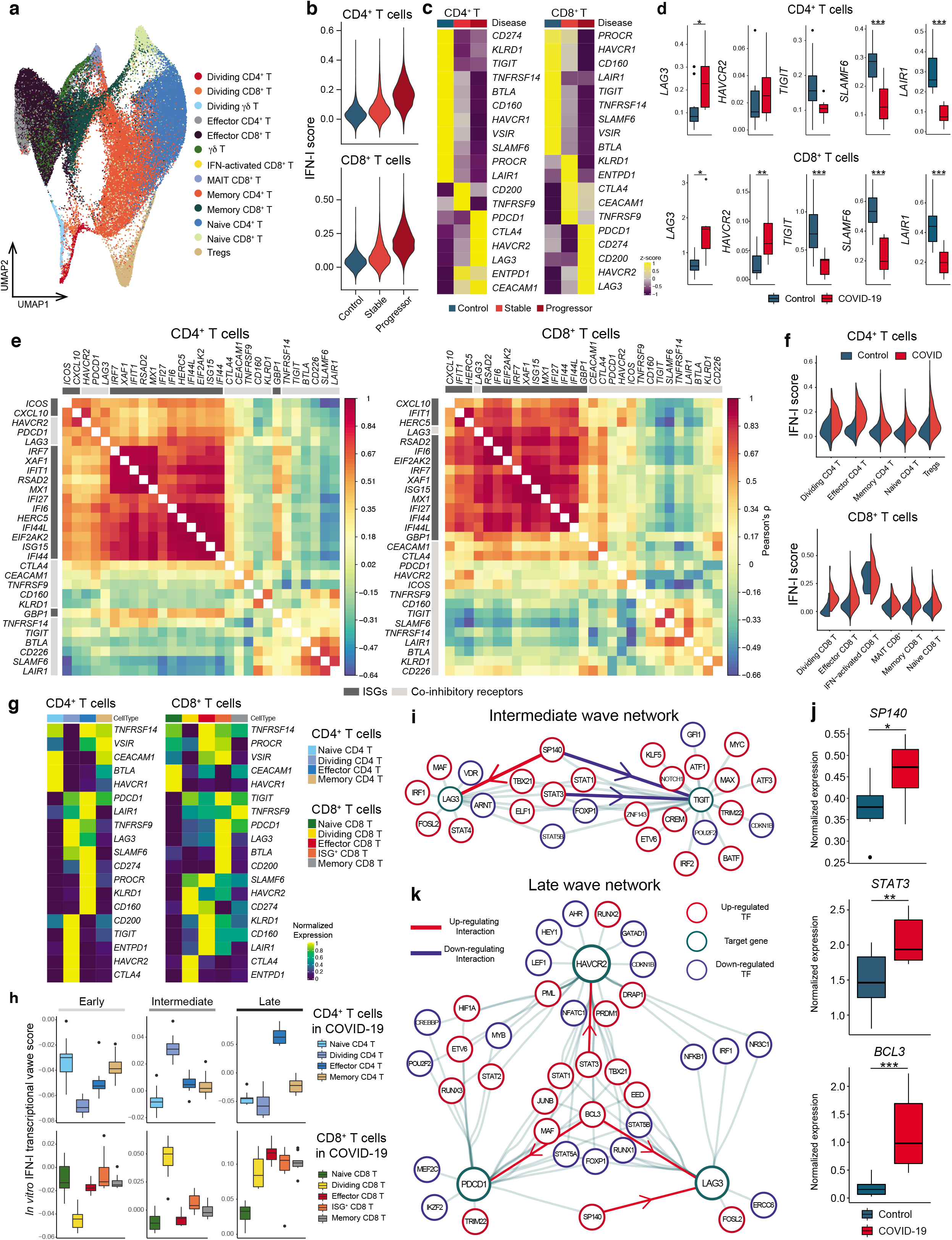
Integration of IFN-I regulatory network with T cells signature in COVID-19. **a,** UMAP representation of T cells from healthy control samples (n = 13) and COVID-19 samples (n = 18). 13 subcluster were identified. **b,** IFN-I score for CD4^+^ and CD8^+^ T cells across the three disease conditions. **c,** Heatmaps for co-inhibitory receptors expression in CD4^+^ and CD8^+^ T cells across the three disease conditions. **d,** Expression of key co-inhibitory receptors between control vs COVID-19 for CD4^+^ and CD8^+^ T cells. Average expression per subject for each gene is shown. *p < 0.05, **p < 0.01, ***p < 0.001. Kruskal-Wallis test. **e,** Correlation matrix of ISGs (dark gray) and co-inhibitory receptors (light gray) in CD4^+^ and CD8^+^ T cells in COVID-19 patients. **f,** IFN-I score for subsets of CD4^+^ and CD8^+^ T cells between control vs COVID-19. **g,** Heatmap showing co-inhibitory receptors expression for subsets of CD4^+^ and CD8^+^ T cells in COVID-19. **h,** Computed three transcriptional waves (early, intermediate, and late) score for the subsets of CD4^+^ and CD8^+^ T cells in COVID-19 patients. Scores were calculated based on upregulated DEGs of CD4^+^ and CD8^+^ T cells for each transcriptional wave. **i,** Regulatory relationship between regulators in intermediate phase network for LAG3 and TIGIT are shown. Positive regulation (TF to target) is highlighted in red and negative regulations in blue. **j,** Box plots showing expression of key regulators between control vs COVID-19 for CD4^+^ T cells. Average expression per subject for each gene is shown. *p < 0.05, **p < 0.01, ***p < 0.001. Kruskal-Wallis test. **k,** Regulatory relationship between regulators in late phase network for *LAG3, HAVCR2,* and *PDCD1* are shown. Positive regulation (TF to target) is highlighted in red.

We then examined which subpopulations were more enriched in the three transcriptional waves of IFN-I response. DEGs specific for each wave were used to compute the scores for the CD4^+^ and CD8^+^ T cell subpopulations in COVID-19 patients (Table 1, Methods). We found that T cells induced *in vitro* with IFN-I strongly mirrored the intermediate wave score on dividing CD4^+^ and CD8^+^ T cells, and the late wave score on effector CD4^+^ T cells and ISG^+^ CD8^+^/effector CD8^+^ T cells (Figure 5h) in COVID-19 patients. Given that expansion of dividing CD4^+^/CD8^+^ T cells are a unique characteristic of COVID-19 patients, we applied the intermediate phase IFN-I regulatory network with the dividing CD4^+^ T cell gene expression signature to examine the relationships between regulators and target genes in this subpopulation. This analysis highlights the regulators that function in establishing the characteristics of the dividing CD4^+^ T cell population under IFN-I response *in vivo* (Supplementary Figure 5d). As LAG-3 is the most upregulated co-inhibitory receptor in the dividing CD4^+^ T cell population, we utilized the network analysis to elucidate the specific regulation of *LAG3* and *TIGIT* that were regulated in an opposite manner under IFN-I response both *in vivo* (Figure 5c, d) and *in vitro* (Figure 1). Analysis of the intermediate wave gene regulatory network demonstrated that *SP140* is a bi-directional regulator for *LAG3* and *TIGIT* under IFN-I response, which is supported by the observation in COVID-19 patients, where elevated *SP140* and *LAG3* but decreased *TIGIT* expression were demonstrated (Figure 5c, d, i, j, Supplementary Figure 5c). In addition, the late wave network demonstrated the complex interaction of regulators for *LAG3, HAVCR2,* and *PDCD1,* in which *BCL3* and *STAT3* are highlighted as validated positive regulators on *LAG3 and HAVCR2* respectively (Figure 5k). Importantly, both *BCL3* and *STAT3* were highly elevated in T cells in COVID-19 patients (Figure 5j). This can be highly relevant to effector T cells development under acute viral infection in which those co-inhibitory receptors play critical role on regulating effector function^33^ and, of note, the late wave signature was enriched in effector T cells in COVID-19 (Figure 5h). These findings strongly suggest that *in vitro* regulatory network can be utilized as a strong tool to explore human acute viral response *in vivo.*

## Discussion

Here, our systematic, computational and biological approach identifies IFN-I as a major driver of co-inhibitory receptor regulation in human T cells. While classical ISG induction has been extensively studied, those investigations have focused primarily on the canonical JAK-STAT pathway downstream of IFN-I receptor. Given that IFN-I exhibits multiple functions in contextdependent roles, a more complex understanding of the IFN-I response beyond this canonical pathway with a more extensive analysis of ISG transcriptional regulation in T cells is critical for elucidating the mechanism of co-inhibitory receptor regulation.

In these studies, we build a dynamic gene regulatory network that controls IFN-I response, and identified key regulatory modules of ISG transcription in T cell responses to IFN-β. Our approach unveiled two mutually antagonistic modules of ISG regulators, which, when acting concordantly, may explain how the harmonized IFN-I induced T cell response is achieved. Within the two modules, we highlighted SP140 as a potential regulator that controls LAG3 and TIGIT in an opposing manner, and STAT3 as a unique positive regulator for TIM-3. These findings provide novel insight into the landscape of the ISG transcriptional network, and sheds light on the large contribution of the noncanonical IFN-I pathway during IFN-I response in T cells^34–36^. Although the newly identified regulators (e.g. *SP140, BCL3)* in this study are not necessarily directly downstream of the conventional JAK/STAT pathway and may act differently depending on the context, they are nevertheless attractive targets for manipulation of specific downstream functional molecules such as co-inhibitory receptors in T cells.

We demonstrate the relevance of our *in vitro* T cell IFN-I response by integrating scRNA-seq from COVID-19 patients, where a predominant T cell IFN-I response was observed. Intriguingly, the expression pattern of co-inhibitory receptors on T cells *in vitro* are highly replicated in severe COVID-19 cases, and classical ISGs were well correlated with one module of co-inhibitory receptors *(LAG3/PDCD1/HAVCR2),* but not with the other modules *(TIGIT/CD160/BTLA/LAIR1).* While dynamics of IFN-I on T cells from COVID-19 patients should be taken into account with higher temporal profiling, we confirmed that the IFN-I response was clearly reduced at a later time point; thus, our data based on earlier collection of blood should reflect the active ISG transcriptomics during human acute viral response. Given the IFN-I response has been shown to contribute to chronic viral infection and cancer, the novel regulators we identified can be examined in these diseases. Further investigations of factors that characterize acute viral infections is likely to differ from chronic viral infections and will be of interest to explore in the context of long-term human infections not well modeled by COVID-19.

In conclusion, our systems biology approach identifies the cytokine signals and regulatory mechanisms that drive expression of co-inhibitory receptors in humans, and provides a pathway to comprehensively capture the dynamics of their expression in humans. Our results will also advance the understanding of the host immune response to a variety of viral infections, and could serve as a resource for mining of existing datasets. Uncovering novel ISG regulators controlling co-inhibitory receptors will create a foundation for further development of new therapeutics for a multitude of different malignant and infectious diseases.

## Figure legends for Supplementary Figures

**Supplementary Figure 1**

**a,** FACS gating strategy for isolating naïve CD4^+^ and naïve CD8^+^ T cells. FACS isolated cells were immediately plated on 96 well round bottom plates coated with anti-CD3 (2 μg/ml) and soluble anti-CD28 (1μg/ml) in the absence or presence of human IL-27 (100 ng/ml) or IFN-β (500 U/ml). **b,** Representative histograms of surface expression of TIM-3, LAG-3, and PD-1 assessed by flow cytometry at 72-96 hours after stimulation. Percent single positive cells for TIM-3, LAG-3, and PD-1 (left) and triple positive cells (right) are shown (n = 6-8). *p < 0.05, **p < 0.01, ****p < 0.0001. Two-way ANOVA or Repeated-measures one-way ANOVA with Tukey’s multiple comparisons test. **c,** Representative dot plots of flow cytometry analysis for TIM-3 and TIGIT expression in naïve CD4^+^ and CD8^+^ T cells (left). Cells were treated as **a,** and analyzed at 72 hours of culture. Percent TIGIT positive cells in naïve CD4^+^ T are shown (n = 8). *p < 0.05, **p < 0.01. Repeated-measures one-way ANOVA with Tukey’s multiple comparisons test (middle). qPCR analysis of *TIGIT* expression over the time course (13 time points from 0 to 96 hours). Each dot represents average expression of two independent individuals’ data (right). ****p < 0.0001. One-way ANOVA with Tukey’s multiple comparisons test. **d,** qPCR analysis of *IL10* and *IFNG* expression over the time course (13 time points from 0 to 96 hours). Each dot represents average expression of two independent individuals’ data (left). IL-10 and IFN-γ production assessed by intracellular staining (right). Cells are treated as in a, and cytokines are stained intracellularly. Cytokine positive cells are detected by flow cytometry (n = 6). *p < 0.05, **p < 0.01. Repeated-measures one-way ANOVA with Tukey’s multiple comparisons test.

**Supplementary Figure 2**

Representative plots for T cell proliferation assay using cell trace violet dye. Naive and memory CD4^+^ T cells were stimulated with anti-CD3 and anti-CD28 in the absence or presence of IFN-β. TIM-3 expression and cellular proliferation were assessed at 24, 48, 72, and 96 hours after stimulation. Overlayed histogram for control and IFN-β condition were shown at right.

**Supplementary Figure 3**

**a,** Schematic experimental setup for high temporal resolution transcriptional profiling. **b,** Heatmap showing log fold change of differentially expressed genes expression between IFN-β and control Th0 condition at each timepoints for naive CD4^+^ (left) and CD8^+^ T cells (right). Genes are clustered based on the three transcriptional wave or bi-modal pattern. **c,** Line plots for *IFI6, IFNG, LAG3,* and *OSM* expression in naive CD4^+^ (left) and CD8^+^ T cells (right).

**Supplementary Figure 4**

**a,** Contour plots for total living cells and backgating analysis for GFP positive cells. Primary naïve CD4^+^ T cells were transduced with scramble shRNA control LV with or without Vpx-VLPs pre-transduction. Cells are collected at 96 hours after starting stimulation and analyzed by flow cytometry. **b, c,** Heatmaps showing the effect of TFs perturbation under IFN-β stimulation on ISGs **(b)** and co-inhibitory receptors **(c)**. Values in the heatmap were normalized by subtractions of log10 fold change of scramble shRNA control over perturbed expression. The “+” sign indicates statistically significant effect with adjusted p value < 0.05 (details in Methods).

**Supplementary Figure 5**

**a, b**, UMAP representation of T cells from healthy control samples (n = 13) and COVID-19 samples (n = 18) color coded by **a**, disease conditions and **b**, each individual. Cells from same individual were labeled as one subject code, which resulted in 10 individual codes shown in **b. c**, Heatmap showing the expression of DETFs for CD4^+^ and CD8^+^ T cells in each T cell subset. **d**, Bundled regulatory network showing interaction between regulators at intermediate phase and transcriptional signature of dividing CD4^+^ T cells in COVID-19. Regulators at intermediate phase are marked with circles (red; upregulated TFs, blue; downregulated TFs), and genes that are differentially expressed in dividing CD4^+^ T cells in COVID-19 were marked with squares (light red; upregulated DEGs, light blue; downregulated DEGs).

## Methods

### Study subjects

Peripheral blood was drawn from healthy controls who were recruited as part of an Institutional Review Board (IRB)-approved study at Yale University, and written consent was obtained. All experiments conformed to the principles set out in the WMA Declaration of Helsinki and the Department of Health and Human Services Belmont Report.

### Human T cell isolation and culture

Peripheral blood mononuclear cells (PBMCs) were prepared from whole blood by Ficoll gradient centrifugation (Lymphoprep, STEMCELL Technologies) and used directly for total T cell enrichment by negative magnetic selection using Easysep magnetic separation kits (STEMCELL Technologies). Cell suspension was stained with anti-CD4 (RPA-T4), anti-CD8 (RPA-T8), anti-CD25 (clone 2A3), anti-CD45RO (UCHL1), anti-CD45RA (HI100) and anti-CD127 (hIL-7R-M21, all from BD Biosciences) for 30 minutes at 4°C. Naïve CD4^+^ T cells (CD4^+^/CD25^neg^/CD127^+^/CD45RO^neg^/CD45RA^+^) and naïve CD8^+^ T cells (CD8^++^/CD25^neg^/CD127^+^/CD45RO^neg^/CD45RA^+^) were sorted on a FACSAria (BD Biosciences). Sorted cells were plated in 96-well round-bottom plates (Corning) and cultured in RPMI 1640 medium supplemented with 5 % Human serum, 2 nM L-glutamine, 5 mM HEPES, and 100 U/ml penicillin, 100 μg/ml streptomycin, 0.5 mM sodium pyruvate, 0.05 mM nonessential amino acids, and 5% human AB serum (Gemini Bio-Products). Cells were seeded (30,000-50,000/wells) into wells pre-coated with anti-human CD3 (2 μg/ml, clone UCHT1, BD Biosciences) along with soluble anti-human CD28 (1 μg/ml, clone 28.2, BD Biosciences) in the presence or absence of human IFN-β (500 U/ml: Pestka Biomedical Laboratories) or IL-27 (100 ng/ml: BioLegend) without adding IL-2.

### Lentiviral and Vpx-VPLs production

Lentiviral plasmids encoding shRNA were obtained from Sigma-Aldrich. Each plasmid was transformed into One Shot Stbl3 chemically competent cells (Invitrogen) and purified by ZymoPURE plasmid Maxiprep kit (Zymo research). Lentiviral pseudoparticles were obtained after plasmid transfection of 293FT cells using Lipofectamine 2000 (Invitrogen) or TurboFectin 8.0 Transfection Reagent (Origene). To prepare Vpx-VLPs, 293T cells were co-transfected by Lipofectamine 2000 or TurboFectin 8.0 Transfection Reagent with the 5 μg pMDL-X, 2.5 μg pcRSV-Rev, 3.5 μg X4-tropic HIV Env, and 1 μg pcVpx/myc, as described previously with some modifications^37,38^. The medium was replaced after 6-12 h with fresh media with 1X Viral boost (Alstem). The lentivirus or Vpx-VLPs containing media was harvested 72 h after transfection and concentrated 80 times using Lenti-X concentrator (Takara Clontech) or Lenti Concentrator (Origene). LV particles were then resuspended in RPMI 1640 media without serum and stored at −80°C before use. Virus titer was determined by using Jurkat T cells and Lenti-X GoStix Plus (Takara Clontech).

### Lentiviral transduction with Vpx-VPLs

Two step Vpx-VLP and LV transduction was performed as described previously with some modiciations^36^. Vpx are pseudotyped with X4-tropic HIV Env to promote efficient entry of Vpx-VLPs into quiescent human T cells^37^. FACS-sorted naïve CD4^+^ T cells were plated at 50,000 cells/well in round bottom 96 well plate and chilled on ice for 15 min. 50 μl of Vpx-VLPs were added to each well and mixed with cold cells for an additional 15 min, then spinfected with highspeed centrifugation (1200 g) for 2 hour at 4 °C. Immediately after centrifugation, cells are cultured overnight at 37 °C. Vpx-transduced cells are spinoculated again with LV particles containing shRNAs with high-speed centrifugation (1000 g) for 1.5 hours at RT. After 24 hours of incubation, the second transduction of LV particles with shRNAs were performed as well as the first time spinoculation. After a second LV transduction, cells were washed and plated into 96-well round-bottom plates pre-coated anti-human CD3 (2 μg/ml) with soluble anti-human CD28 (1 μg/ml), in the presence or absence of human IFN-β (500 U/ml). Cells are collected at day 4 after anti-CD3/CD28 stimulation and GFP positive cells were sorted by FACSAria.

### Real time quantitative PCR

Total RNA was extracted using RNeasy Micro Kit (QIAGEN), or ZR-96 Quick-RNA kit (Zymo Research), according to the manufacturer’s instructions. RNA was treated with DNase and reverse transcribed using TaqMan Reverse Transcription Reagents (Applied Biosystems) or SuperScript IV VILO Master Mix (Invitrogen). cDNAs were amplified with Taqman probes (Taqman Gene Expression Arrays) and TaqMan Fast Advanced Master Mix on a StepOne Real-Time PCR System (Applied Biosystems) according to the manufacturer’s instructions. Relative mRNA expression was evaluated after normalization with *B2M* expression.

### Flow cytometry analysis

Cells were stained with LIVE/DEAD Fixable Near-IR Dead Cell Stain kit (Invitrogen) and surface antibodies for 30 min at 4 °C. For intracellular cytokine staining, cells were treated with 50 nM phorbol-12-myristate-13-acetate (MilliporeSigma) and 250 nM ionomycin (MilliporeSigma) for 4 hours in the presence of Brefeldin A (BD Biosciences) before harvesting. Cells were washed and fixed with BD Cytofix™ Fixation Buffer (BD Biosciences) for 10 min at RT, then washed with PBS. Intracellular cytokines were stained in permeabilization buffer (eBioscience) for 30 min at 4 °C. The following antibodies were used: anti-LAG-3 (11C3C65, BioLegend), anti-PD-1 (EH12.1, BD Biosciences), anti-TIGIT (MBSA43, eBioscience), anti-Tim-3 (F38-2E2, Biolegend), anti-IFN-γ (4S.B3, eBioscience), and IL-10 (JES3-9D7, Biolegend). Cells were acquired on a BD Fortessa flow cytometer and data was analyzed with FlowJo software v10 (Threestar).

### RNA-seq library preparation and data analysis

cDNAs were generated from isolated RNAs using SMART-Seq v4 Ultra Low Input RNA Kit for sequencing (Takara/Clontech). Barcoded libraries were generated by the Nextera XT DNA Library Preparation kit (Illumina) and sequenced with a 2×100 bp paired-end protocol on the HiSeq 4000 Sequencing System (Illumina).

After sequencing, adapter sequences and poor-quality bases (quality score < 3) were trimmed with Trimmomatic. Remaining bases were trimmed if their average quality score in a 4 bp sliding window fell below 5. FastQC was used to obtain quality control metrics before and after trimming. Remaining reads were aligned to the GRCh38 human genome with STAR 2.5.2^39^. We used Picard to remove optical duplicates and to compile alignment summary statistics and RNA-seq summary statistics. After alignment, reads were quantitated to gene level with RSEM^40^ using the Ensembl annotation.

### Identification of three transcriptional waves

The correlation matrix is created by Pearson correlating^41^ the IFN-β expression profile of each time point with all the other time points, creating a symmetric matrix of Pearson correlation coefficients.

### Differential expression calculation

The differential expression (DE)^42^ of the sequenced genes from every time point was calculated. The DE was calculated using the DEseq2^43^ R package. An in-house decision algorithm was built to determine which genes are DE. The algorithm used three separate testing methods available in DEseq2 for calculating DE genes: Wald^44^, likelihood ratio test (LRT)^45^, and timecourse^46^. For each of the three methods, genes with false discovery rate (FDR) adjusted P.value bellow 0.05, are regarded as DE. The algorithm defined genes as DE differently for CD4^+^ and CD8^+^ T cells. For CD4^+^ T cells, if TFs appeared in two out of the three calculating methods (agree by two), they were regarded as DE. For CD8^+^ T cells, if TFs appeared in any of the methods above, they were regarded as DE.

### Selection of regulators for perturbation

The list of TFs for perturbation was selected based on following aspects: 1) overlapped differentially expressed TFs across the time point between CD4^+^ and CD8^+^ T cells in our results. Intersection of DETFs in our in vitro data were chosen; 2) differentially expressed TFs in human tumor infiltrated T cells^21–24^ that were significantly correlated with exhausted T cell cluster where LAG-3/PD-1/TIM-3 were highly upregulated. ‘Human TIL score’ for each gene was calculated by the number of times it was shared between the four different human cancer TIL datasets^21–24^; 3) HIV specific T cell signature in progressive patients compared to stable patients^25^. 4) TFs that were included by IL-27 and categorized as IL-27 driven co-inhibitory receptor modules^3^. ‘Human ISG score’ for each gene was calculated by the number of times it was shared between the three different categories (T cells, PBMCs, and all immune cells) of human ISGs identified by Interferome database. All perturbed TFs were confirmed as IFN-I responsible genes that showed ‘human ISG score’ more than 1.

### Heatmap of perturbed TFs

21 TFs were perturbed using lentiviral shRNA, together with a scramble shRNA control (SCR). In order to better understand the effect of perturbing said TFs, selected genes of interest (GOIs) were analyzed. The calculation of the heatmap values in Supplementary Figure 4b and c is as follows: first the expression values of perturbed GOIs is divided by control values without perturbation (fold change), this quotient is then logged in base 10. The result of the logarithm of the GOIs is the subtracted by the logarithm of SCR fold change.

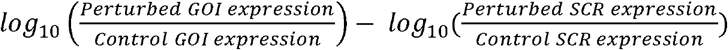

DE analysis was conducted for perturbation against control, genes who yielded n FDR adjusted P.value lower than 0.05 were regarded as significant and display a white plus on their tile.

### PCA and PCA biplot analysis

The PCA was conducted on a DEGs defined as above as variables, and perturbed genes as observations. The data was normalized by the scramble shRNA control identically to the Heatmap of perturbed TFs. Although *PDCD1* was not defined as DE genes across the time course, it was clearly differentially expressed at later time points in mRNA-seq and qPCR, thus we manually depicted in Figure 5f. Genes of IFN Score B^47^ (see Table 1) were represented as ISGs in Figure 3e. The PCA and biplot analysis were calculated and visualized using the R package FactoMineR^48^.

### Regulatory networks

Following DE analysis in each transcriptional wave, the DE genes were separated to TFs and their targets (from->to). The targets of all DETFs were determined using ChIP-seq data from the database GTRD^49^. TFs and targets defined as DE were added as network nodes, and edges (connections) were added between them. The network figures were created using the software Cytoscape^50^.

### Top Regulatory TFs Heatmaps

We ranked the DE TFs of each transcriptional time wave to identify which are the most dominant in the overall differentiation process. HG stands for hyper-geometric, the value in the heatmap is the-*log*_10_(*P.value*) of a hypergeometric enrichment test. The targets of each TF are tested for enrichment of DE targets in the network, relatively to targets that aren’t DE in the network. The HG calculation was conducted using the python SciPy package^51^. Cent stands for centrality, which is a parameter that is given to each node, based on the shortest path from the node to the other nodes in the network. It represents how central and connected a node is in the rest of the network^52^. The centrality calculation was conducted using the python NetworkX^53^ package. The rank column is an average of both HG and Cent values, after normalization.

### Integration of Perturbation Data to Regulatory Networks

DE analysis was conducted for perturbation against control. Genes that were significantly affected by a TF perturbation were added as a “validated” edges between the perturbed TF and the target gene. If a gene was up-regulated by a TF perturbation, the interaction between them is registered as down-regulation. If a gene was down-regulated by a TF perturbation, the interaction between them is registered as up-regulation.

### Backbone Hierarchical Networks

Using the software Cytoscape, we implemented a hierarchical layout, which takes into account the directionality of the connections between the TFs. A TF which has only outgoing connections will be placed at the top of the hierarchy, while a TF which has only incoming connections will be placed at the bottom.

### Bridging TFs Network

DETFs and their targets from the three transcriptional waves were combined to create a comprehensive network of the dynamic between transcriptional waves. TFs and their targets were annotated by the transcriptional wave in which they are DE. TFs that appear in more than one transcriptional wave are regarded as bridging TFs.

### Reanalysis of COVID-19 single-cell RNA sequencing data

A PBMC single cell RNA seq data set of 10 COVID-19 patients and 13 matched controls was reanalyzed which had been previously performed and reported by us^32^. We have described the full cohort and detailed methods elsewhere^32^. From eight of the ten COVID-19 samples, PBMCs from two different time points had been analyzed. Four of the COVID-19 patients had been classified as progressive, the other six COVID-19 patients as stable. Informed consent had been obtained of all subjects and the protocol had been approved by Yale Human Research Protection Program Institutional Review Boards (FWA00002571, Protocol ID. 2000027690). Briefly, single cell barcoding of PBMCs and library construction had been performed using the 10x Chromium NextGEM 5prime kit according to manufacturer’s instructions. Libraries had been sequenced on an Illumina Novaseq 6000 platform. Raw reads had been demultiplexed and processed using Cell Ranger (v3.1) mapping to the GRCh38 (Ensembl 93) reference genome. Resulting gene-cell matrices had been analyzed using the package Seurat^54^ in the software R^55^ including integration of data, clustering, multiplet identification and cell type annotation. The final annotated R object was used and re-analyzed in Seurat with default settings – unless otherwise specified – as follows:

The three cell populations “Dividing T & NK”, “Effector T” and “Memory CD4 & MAIT” were each subsetted and reclustered to obtain a finer cell type granularity as they included a mix of CD4, CD8, MAIT and gamma delta T cells. Per subset, the top 500 variable genes were determined by the “FindVariableFeatures” function using the “vst” method. Data was scaled using the “ScaleData” function regressing out the total number of UMI and the percentage of UMIs arising from the mitochondrial genome. After Principal Component (PC) Analysis, the first 10 Principal Components (PCs) were utilized to detect the nearest neighbors using the “FindNeighbors” function and clustered by Seurat’s Louvain algorithm implementation “FindClusters” using a resolution of 0.2 for “Dividing T & NK”, of 0.3 for “Effector T” and of 0.1 for “Memory CD4 & MAIT” subsets. Cluster-specific gene expression profiles were established using the “FindAllMarkers” per cluster and per subset to annotate the clusters. New cell type annotations were then transferred back to the full dataset.

A new Uniform Approximation and Projection (UMAP) embedding was created by integrating the datasets on a subject level as follows: A subset containing all T cells was generated, which was then split by subject. For each subject, the top 2000 variable genes were selected, then integration anchors determined by “FindIntegrationAnchors” (with k.filter = 150). These anchors were used to integrate the data using the “IntegrateData” function with top 30 dimensions. The integrated data was scaled, subjected to a PC analysis and the top 13 PCs used as input for the “RunUMAP” function on 75 nearest neighbors^56^.

Module scores were calculated using the “AddModuleScore” function using a) all genes within the GO list “RESPONSE TO TYPE I INTERFERON” (GO:0034340)^57^ and b) all genes significantly associated with either of the three waves in our *in vitro* perturbation experiments (see Table 1). Differential gene expression was established using Seurat’s implementation of the Wilcoxon Rank Sum test within the “FindMarkers” function with a Bonferroni correction for multiple testing.

### Statistical analysis

Detailed information about statistical analysis, including tests and values used, is provided in the figure legends. P-values of 0.05 or less were considered significant.

### Data and software availability

The sequence data generated in this study will be deposited in the Gene Expression Omnibus (GEO) and the accession code will be provided prior to publication.

## Acknowledgements

We thank L. Devine and C. Wang for assistance with FACS based cell sorting; G. Wang and C. Castaldi at Yale Center for Genome Analysis for support with 10x Genomics library preparation and sequencing; K. Raddassi and L. Zhang for processing scRNA-seq samples; L. Geng and M. Zhang for preparation of the bulk RNA-seq libraries, and sequencing; M. Taura for assistance developing Vpx-VLP based lenti viral transduction; N. Chihara and H. Asashima for useful discussions and input on the manuscript.

## Contributions

T.S.S., A.M., V.K.K., and D.A.H. conceptualized the study. T.S.S. performed *in vitro* experiments with the help of H.A.S., M.C., and P-P.A.; T.S.S. prepared sequencing libraries and M.R.L. processed bulk RNA-seq data alignment; S.D. and A.M. performed computational analysis to construct gene regulatory network with input from T.S.S.; J.C.S. performed scRNA-seq analysis on COVID-19 data with the help of T.S.S., A.U., and N.K.; T.S.S., S.D., J.C.S., A.M., and D.A.H. wrote the manuscript with input from all authors; T.S.S., A.M., V.K.K. and D.A.H. supervised the overall study.

## References

1 Wang, Y. et al. Timing and magnitude of type I interferon responses by distinct sensors impact CD8 T cell exhaustion and chronic viral infection. Cell Host Microbe 11, 631–642, doi:10.1016/j.chom.2012.05.003 (2012).

2 McLane, L. M., Abdel-Hakeem, M. S. & Wherry, E. J. CD8 T Cell Exhaustion During Chronic Viral Infection and Cancer. Annu Rev Immunol 37, 457–495, doi:10.1146/annurev-immunol-041015-055318 (2019).

3 Chihara, N. et al. Induction and transcriptional regulation of the co-inhibitory gene module in T cells. Nature 558, 454–459, doi:10.1038/s41586-018-0206-z (2018).

4 DeLong, J. H. et al. IL-27 and TCR Stimulation Promote T Cell Expression of Multiple Inhibitory Receptors. Immunohorizons 3, 13–25, doi:10.4049/immunohorizons.1800083 (2019).

5 Gonzalez-Navajas, J. M., Lee, J., David, M. & Raz, E. Immunomodulatory functions of type I interferons. Nat Rev Immunol 12, 125–135, doi:10.1038/nri3133 (2012).

6 Crow, M. K. & Ronnblom, L. Type I interferons in host defence and inflammatory diseases. Lupus Sci Med 6, e000336, doi:10.1136/lupus-2019-000336 (2019).

7 Musella, M., Manic, G., De Maria, R., Vitale, I. & Sistigu, A. Type-I-interferons in infection and cancer: Unanticipated dynamics with therapeutic implications. Oncoimmunology 6, e1314424, doi:10.1080/2162402X.2017.1314424 (2017).

8 Welsh, R. M., Bahl, K., Marshall, H. D. & Urban, S. L. Type 1 interferons and antiviral CD8 T-cell responses. PLoS Pathog 8, e1002352, doi:10.1371/journal.ppat.1002352 (2012).

9 Axtell, R. C., Raman, C. & Steinman, L. Type I interferons: beneficial in Th1 and detrimental in Th17 autoimmunity. Clin Rev Allergy Immunol 44, 114–120, doi:10.1007/s12016-011-8296-5 (2013).

10 Teijaro, J. R. et al. Persistent LCMV infection is controlled by blockade of type I interferon signaling. Science 340, 207–211, doi:10.1126/science.1235214 (2013).

11 Wilson, E. B. et al. Blockade of chronic type I interferon signaling to control persistent LCMV infection. Science 340, 202–207, doi:10.1126/science.1235208 (2013).

12 Baumeister, S. H., Freeman, G. J., Dranoff, G. & Sharpe, A. H. Coinhibitory Pathways in Immunotherapy for Cancer. Annu Rev Immunol 34, 539–573, doi:10.1146/annurev-immunol-032414-112049 (2016).

13 Anderson, A. C., Joller, N. & Kuchroo, V. K. Lag-3, Tim-3, and TIGIT: Co-inhibitory Receptors with Specialized Functions in Immune Regulation. Immunity 44, 989–1004, doi:10.1016/j.immuni.2016.05.001 (2016).

14 Das, M., Zhu, C. & Kuchroo, V. K. Tim-3 and its role in regulating anti-tumor immunity. Immunol Rev 276, 97–111, doi:10.1111/imr.12520 (2017).

15 Crawford, A. et al. Molecular and transcriptional basis of CD4(+) T cell dysfunction during chronic infection. Immunity 40, 289–302, doi:10.1016/j.immuni.2014.01.005 (2014).

16 Zhen, A. et al. Targeting type I interferon-mediated activation restores immune function in chronic HIV infection. J Clin Invest 127, 260–268, doi:10.1172/JCI89488 (2017).

17 Molle, C. et al. IL-27 synthesis induced by TLR ligation critically depends on IFN regulatory factor 3. J Immunol 178, 7607–7615, doi:10.4049/jimmunol.178.12.7607 (2007).

18 Petricoin, E. F. 3rd, et al. Antiproliferative action of interferon-alpha requires components of T-cell-receptor signalling. Nature 390, 629–632, doi:10.1038/37648 (1997).

19 Marshall, H. D., Urban, S. L. & Welsh, R. M. Virus-induced transient immune suppression and the inhibition of T cell proliferation by type I interferon. J Virol 85, 5929–5939, doi:10.1128/JVI.02516-10 (2011).

20 Son, H. J. et al. Oncostatin M Suppresses Activation of IL-17/Th17 via SOCS3 Regulation in CD4+ T Cells. J Immunol 198, 1484–1491, doi:10.4049/jimmunol.1502314 (2017).

21 Luoma, A. M. et al. Molecular Pathways of Colon Inflammation Induced by Cancer Immunotherapy. Cell 182, 655–671e622, doi:10.1016/j.cell.2020.06.001 (2020).

22 Sade-Feldman, M. et al. Defining T Cell States Associated with Response to Checkpoint Immunotherapy in Melanoma. Cell 175, 998–1013e1020, doi:10.1016/j.cell.2018.10.038 (2018).

23 Oh, D. Y. et al. Intratumoral CD4(+) T Cells Mediate Anti-tumor Cytotoxicity in Human Bladder Cancer. Cell 181, 1612–1625 e1613, doi:10.1016/j.cell.2020.05.017 (2020).

24 Li, H. et al. Dysfunctional CD8 T Cells Form a Proliferative, Dynamically Regulated Compartment within Human Melanoma. Cell 176, 775–789e718, doi:10.1016/j.cell.2018.11.043 (2019).

25 Quigley, M. et al. Transcriptional analysis of HIV-specific CD8+ T cells shows that PD-1 inhibits T cell function by upregulating BATF. Nat Med 16, 1147–1151,doi:10.1038/nm.2232 (2010).

26 Rusinova, I. et al. Interferome v2.0: an updated database of annotated interferon-regulated genes. Nucleic Acids Res 41, D1040–1046, doi:10.1093/nar/gks1215 (2013).

27 Laguette, N. et al. SAMHD1 is the dendritic-and myeloid-cell-specific HIV-1 restriction factor counteracted by Vpx. Nature 474, 654–657, doi:10.1038/nature10117 (2011).

28 Baldauf, H. M. et al. SAMHD1 restricts HIV-1 infection in resting CD4(+) T cells. Nat Med 18, 1682–1687, doi:10.1038/nm.2964 (2012).

29 Chandramohan, V. et al. c-Myc represses FOXO3a-mediated transcription of the gene encoding the p27(Kip1) cyclin dependent kinase inhibitor. J Cell Biochem 104, 2091–2106, doi:10.1002/jcb.21765 (2008).

30 Ptaschinski, C. et al. RSV-Induced H3K4 Demethylase KDM5B Leads to Regulation of Dendritic Cell-Derived Innate Cytokines and Exacerbates Pathogenesis In Vivo. PLoS Pathog 11, e1004978, doi:10.1371/journal.ppat.1004978 (2015).

31 Chen, Z. et al. <em>In vivo</em> CRISPR screening identifies Fli1 as a transcriptional safeguard that restrains effector CD8 T cell differentiation during infection and cancer. bioRxiv, 2020.2005.2020.087379, doi:10.1101/2020.05.20.087379 (2020).

32 Unterman, A. et al. Single-Cell Omics Reveals Dyssynchrony of the Innate and Adaptive Immune System in Progressive COVID-19. medRxiv, 2020.2007.2016.20153437, doi:10.1101/2020.07.16.20153437 (2020).

33 Grosso, J. F. et al. LAG-3 regulates CD8+ T cell accumulation and effector function in murine self-and tumor-tolerance systems. J Clin Invest 117, 3383–3392, doi:10.1172/JCI31184 (2007).

34 Gil, M. P. et al. Biologic consequences of Stat1-independent IFN signaling. Proc Natl Acad Sci U S A 98, 6680–6685, doi:10.1073/pnas.111163898 (2001).

35 Wang, W., Xu, L., Su, J., Peppelenbosch, M. P. & Pan, Q. Transcriptional Regulation of Antiviral Interferon-Stimulated Genes. Trends Microbiol 25, 573–584, doi:10.1016/j.tim.2017.01.001 (2017).

36 Mostafavi, S. et al. Parsing the Interferon Transcriptional Network and Its Disease Associations. Cell 164, 564–578, doi:10.1016/j.cell.2015.12.032 (2016).

37 Geng, X., Doitsh, G., Yang, Z., Galloway, N. L. & Greene, W. C. Efficient delivery of lentiviral vectors into resting human CD4 T cells. Gene Ther 21, 444–449, doi:10.1038/gt.2014.5 (2014).

38 Norton, T. D., Miller, E. A., Bhardwaj, N. & Landau, N. R. Vpx-containing dendritic cell vaccine induces CTLs and reactivates latent HIV-1 in vitro. Gene Ther 22, 227–236, doi:10.1038/gt.2014.117 (2015).

39 Dobin, A. et al. STAR: ultrafast universal RNA-seq aligner. Bioinformatics 29, 15–21, doi:10.1093/bioinformatics/bts635 (2013).

40 Li, B. & Dewey, C. N. RSEM: accurate transcript quantification from RNA-Seq data with or without a reference genome. BMC Bioinformatics 12, 323, doi:10.1186/1471-2105-12-323 (2011).

41 Ly, A., Marsman, M. & Wagenmakers, E. J. Analytic posteriors for Pearson’s correlation coefficient. Stat Neerl 72, 4–13, doi:10.1111/stan.12111 (2018).

42 Bojnordi, M. N. et al. Differentiation of Spermatogonia Stem Cells into Functional Mature Neurons Characterized with Differential Gene Expression. Mol Neurobiol 54, 5676–5682, doi:10.1007/s12035-016-0097-7 (2017).

43 Love, M. I., Huber, W. & Anders, S. Moderated estimation of fold change and dispersion for RNA-seq data with DESeq2. Genome Biol 15, 550, doi:10.1186/s13059-014-0550-8 (2014).

44 Chen, Y. M., Weng, Y. T., Dong, X. & Tsong, Y. Wald tests for variance-adjusted equivalence assessment with normal endpoints. J Biopharm Stat 27, 308–316, doi:10.1080/10543406.2016.1265542 (2017).

45 Rubinstein, M. L., Kraft, C. S. & Parrott, J. S. Determining qualitative effect size ratings using a likelihood ratio scatter matrix in diagnostic test accuracy systematic reviews. Diagnosis (Berl) 5, 205–214, doi:10.1515/dx-2018-0061 (2018).

46 Leong, H. S. et al. A global non-coding RNA system modulates fission yeast protein levels in response to stress. Nat Commun 5, 3947, doi:10.1038/ncomms4947 (2014).

47 El-Sherbiny, Y. M. et al. A novel two-score system for interferon status segregates autoimmune diseases and correlates with clinical features. Sci Rep 8, 5793, doi:10.1038/s41598-018-24198-1 (2018).

48 Lê, S., Josse, J. & Husson, F. FactoMineR: An R Package for Multivariate Analysis. 2008 25, 18, doi:10.18637/jss.v025.i01 (2008).

49 Yevshin, I., Sharipov, R., Kolmykov, S., Kondrakhin, Y. & Kolpakov, F. GTRD: a database on gene transcription regulation-2019 update. Nucleic Acids Res 47, D100–D105, doi:10.1093/nar/gky1128 (2019).

50 Shannon, P. et al. Cytoscape: a software environment for integrated models of biomolecular interaction networks. Genome Res 13, 2498–2504, doi:10.1101/gr.1239303 (2003).

51 Virtanen, P. et al. SciPy 1.0: fundamental algorithms for scientific computing in Python. Nat Methods 17, 261–272, doi:10.1038/s41592-019-0686-2 (2020).

52 Piraveenan, M., Prokopenko, M. & Hossain, L. Percolation centrality: quantifying graph-theoretic impact of nodes during percolation in networks. PLoS One 8, e53095, doi:10.1371/journal.pone.0053095 (2013).

53 Hagberg, A., Swart, P. & Chult, D. Exploring Network Structure, Dynamics, and Function Using NetworkX. (2008).

54 Stuart, T. et al. Comprehensive Integration of Single-Cell Data. Cell 177, 1888–1902.e1821, doi:10.1016/j.cell.2019.05.031 (2019).

55 Team, R. C. (2020).

56 McInnes, L. & Healy, J. UMAP: Uniform Manifold Approximation and Projection for Dimension Reduction. ArXiv abs/1802.03426(2018).

57 Ashburner, M. et al. Gene ontology: tool for the unification of biology. The Gene Ontology Consortium. Nat Genet 25, 25–29, doi:10.1038/75556 (2000).

