## Supplementary Figure1-5 for "Type I Interferon Transcriptional Network Regulates Expression of Coinhibitory Receptors in Human T cells"

Supplementary Figure 1

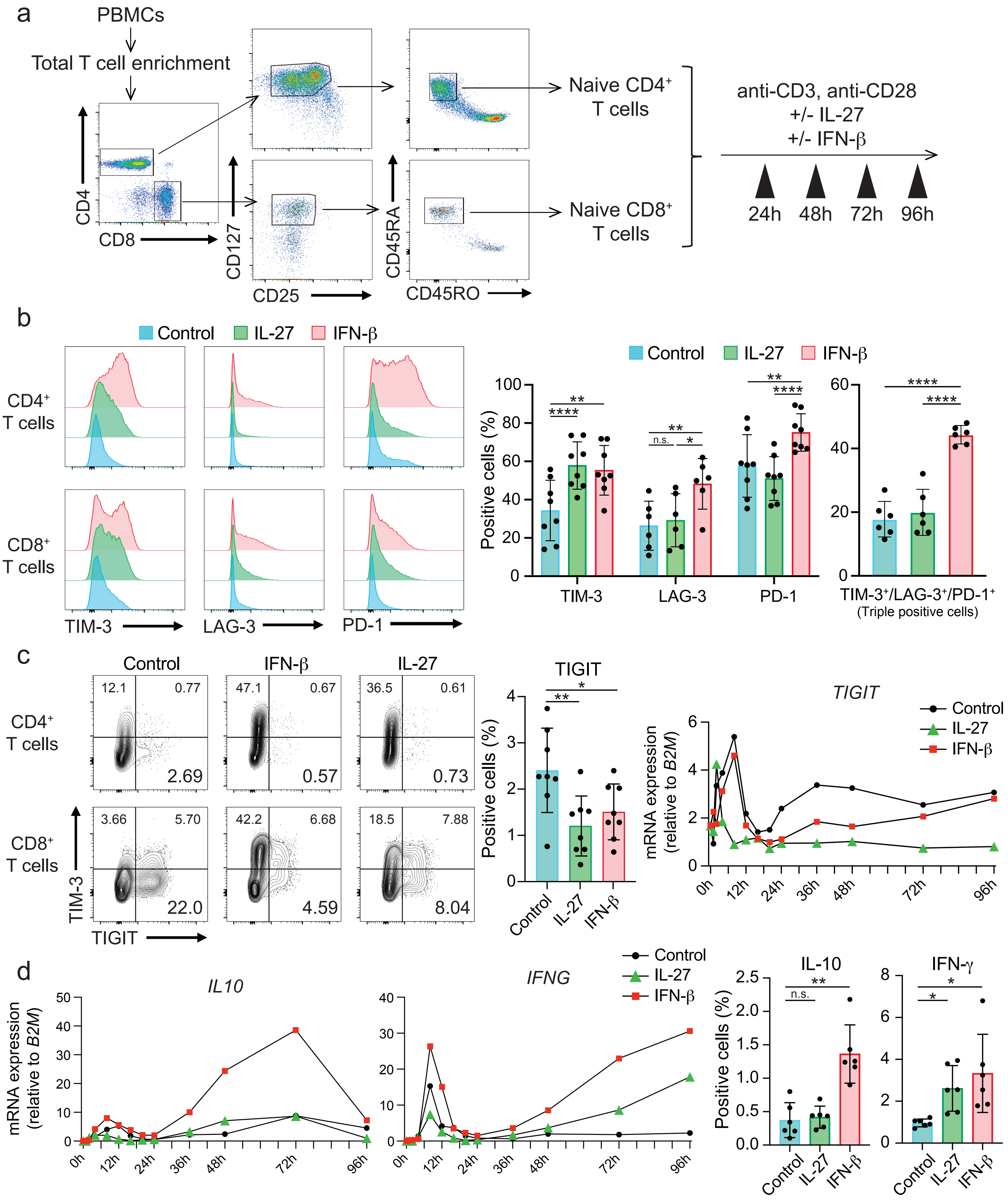

Supplementary Figure 2

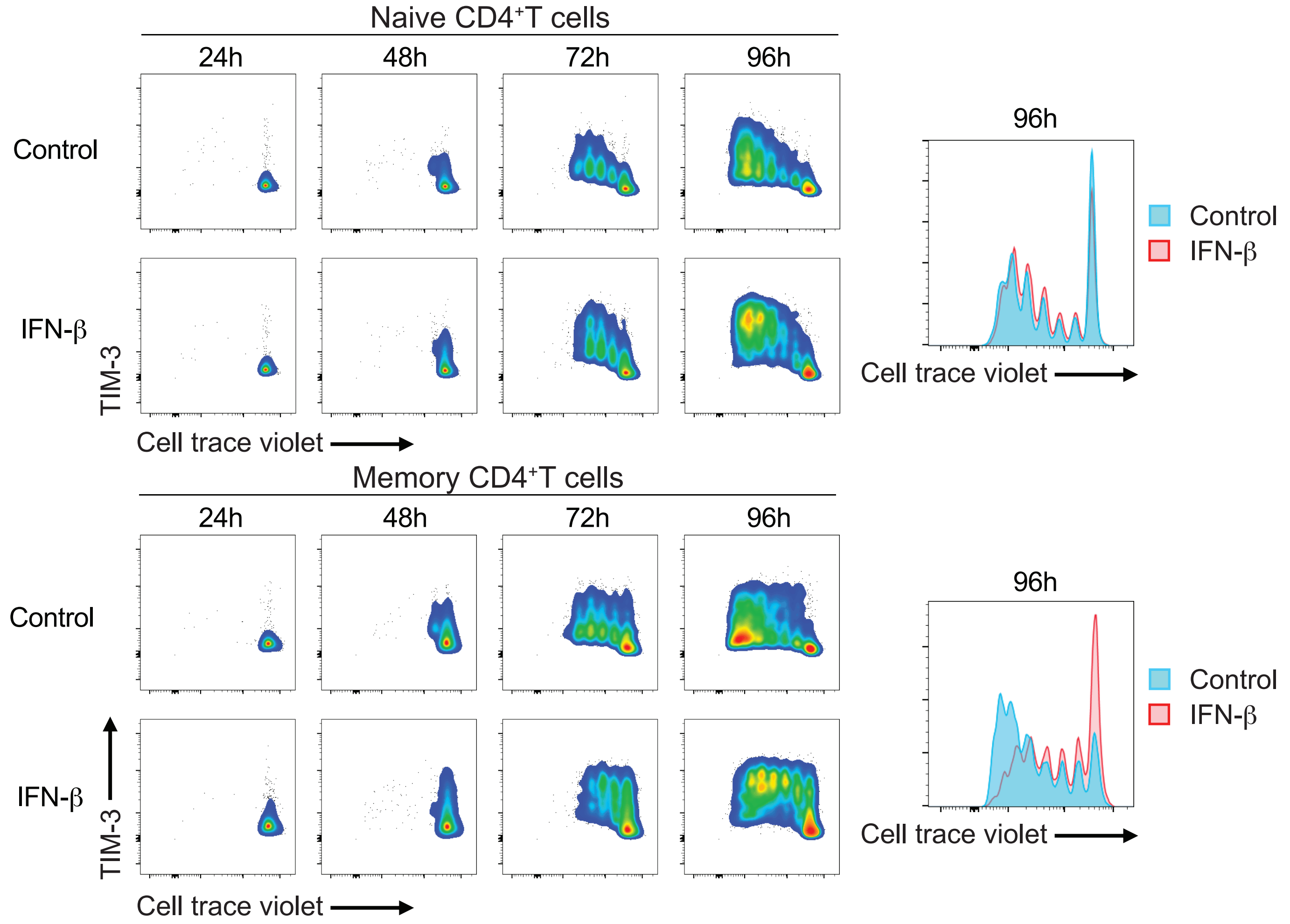

### Supplementary Figure 3

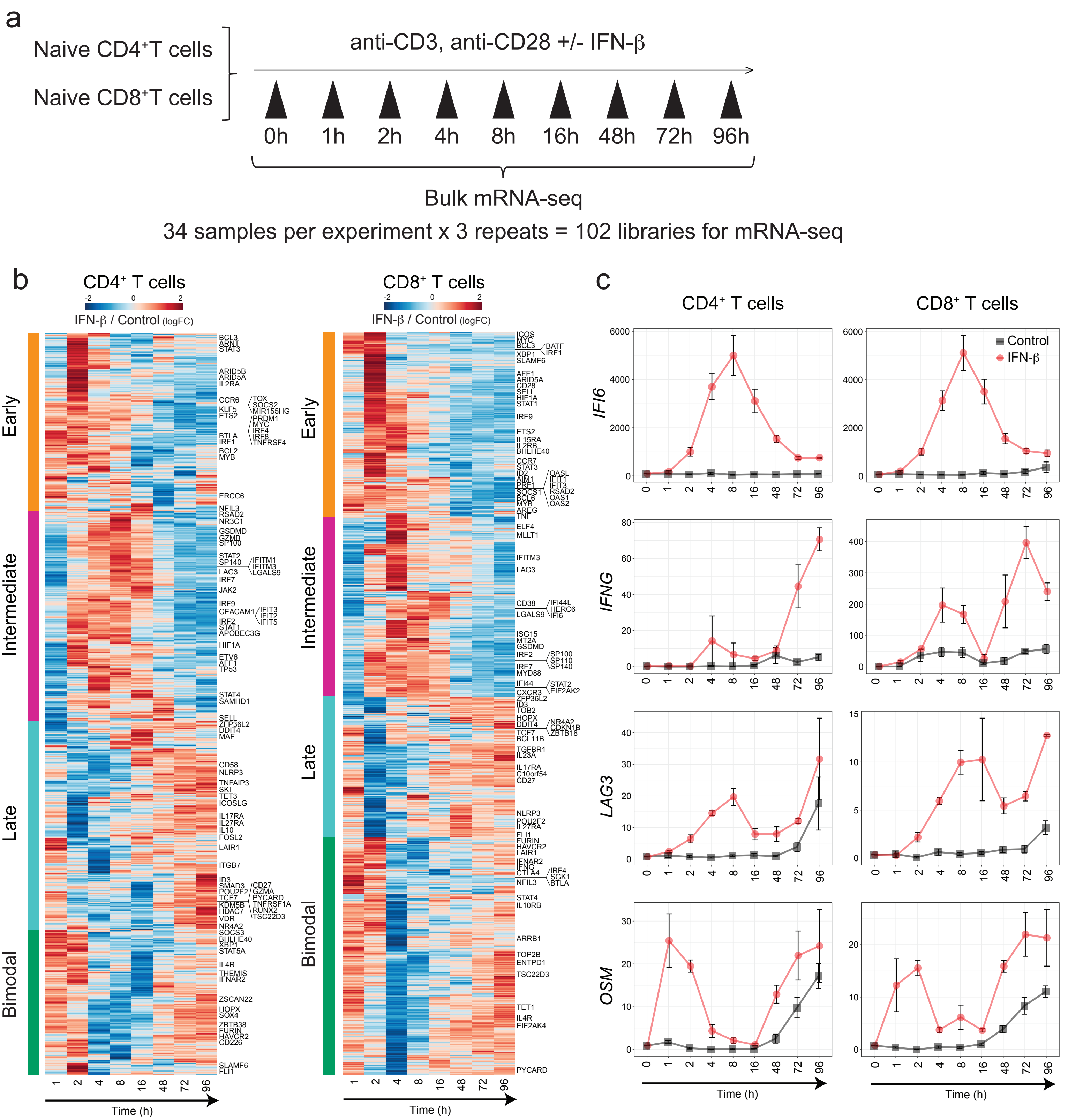

Supplementary Figure 4

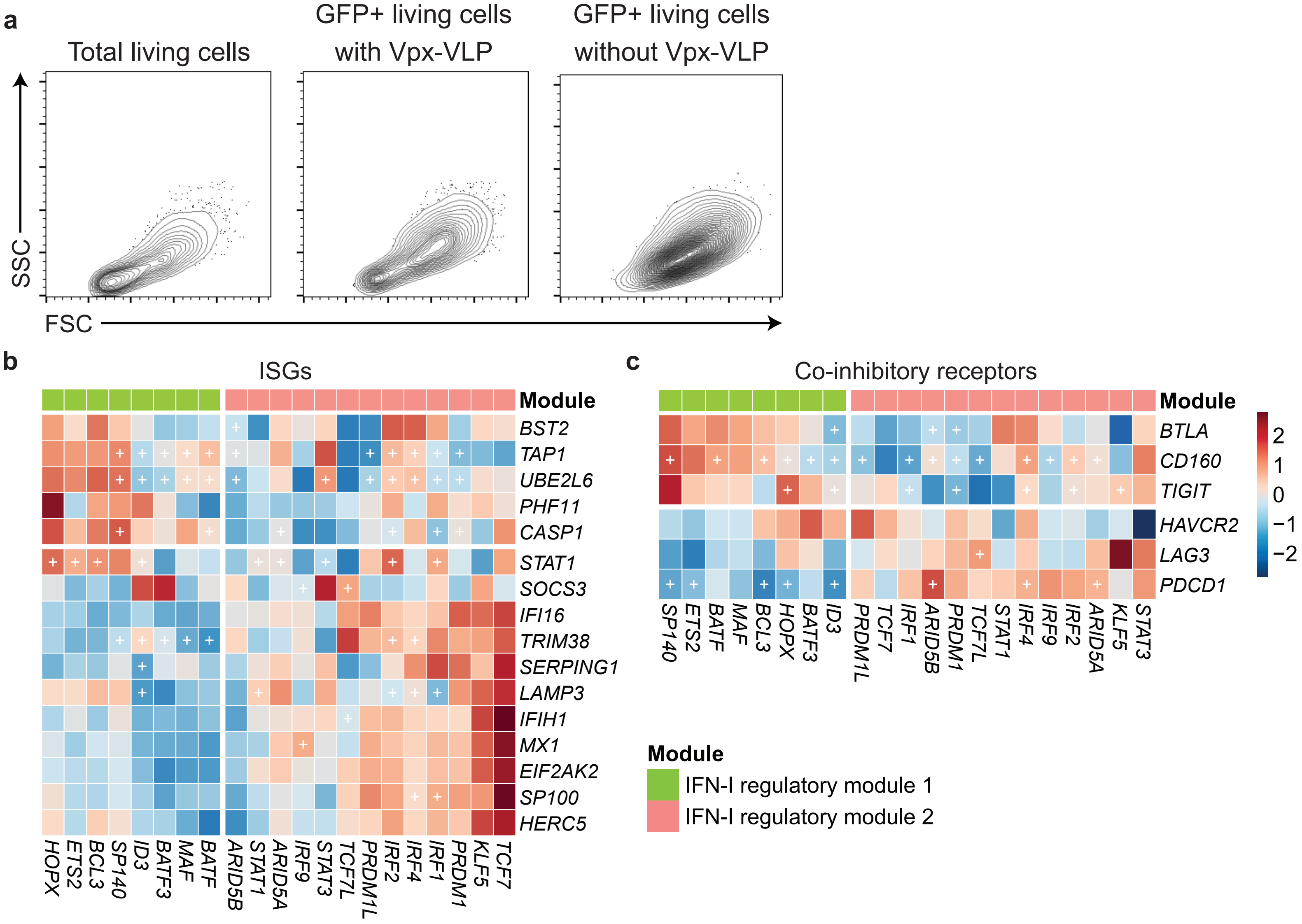

Supplementary Figure 5

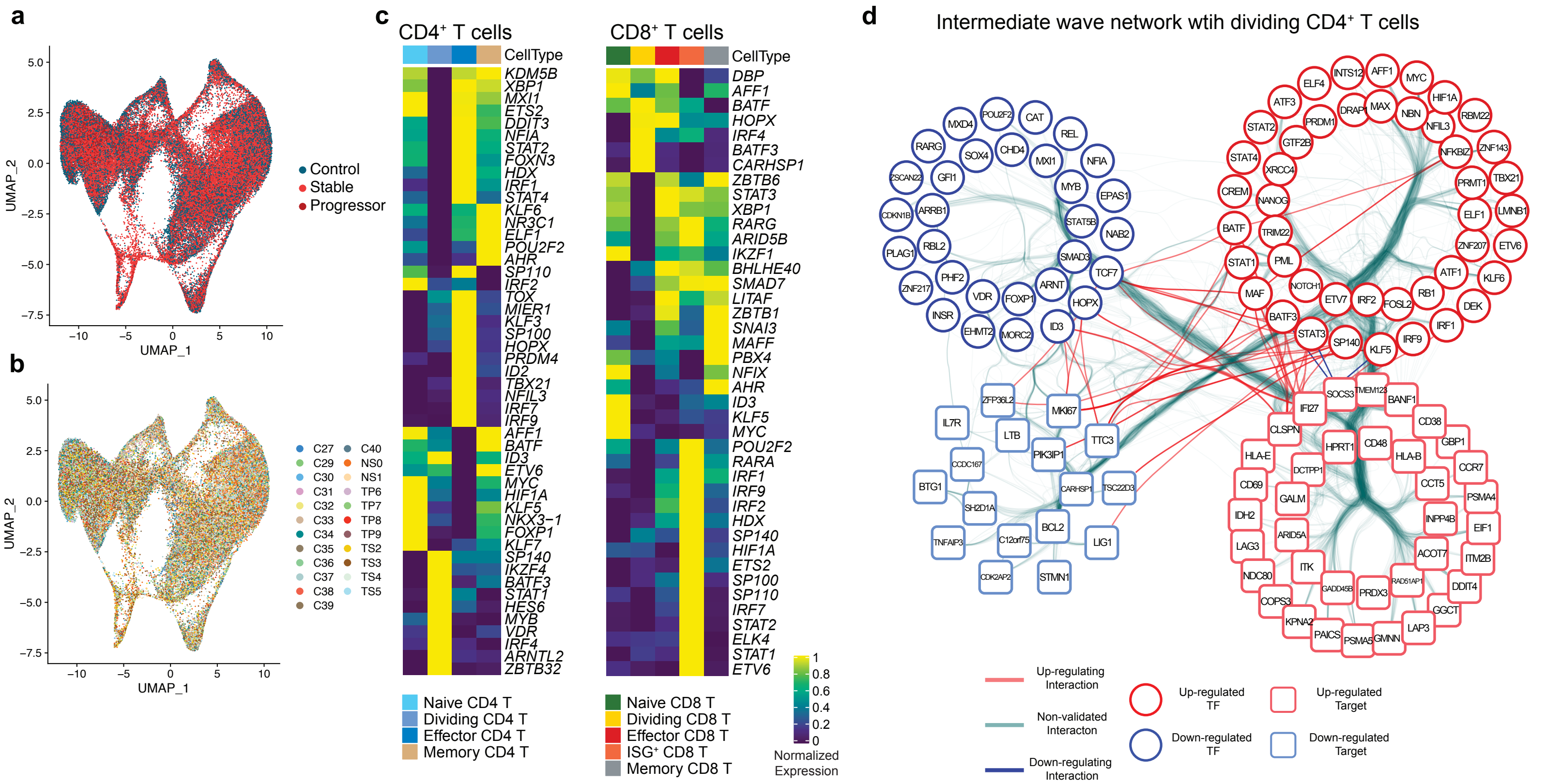
